# Formation of a stable RNase Y-RicT (YaaT) complex requires RicA (YmcA) and RicF (YlbF)

**DOI:** 10.1101/2023.05.22.541740

**Authors:** Eugenie Dubnau, Micaela DeSantis, David Dubnau

## Abstract

In *Bacillus subtilis*, the RicT (YaaT), RicA (YmcA) and RicF (YlbF) proteins, which form a stable ternary complex, are needed together with RNase Y (Rny), to cleave and thereby stabilize several key transcripts encoding enzymes of intermediary metabolism. We show here that RicT, but not RicA or RicF, forms a stable complex with Rny, and that this association requires the presence of RicA and RicF. We propose that RicT is handed off from the ternary complex to Rny. We show further that the two iron-sulfur clusters carried by the ternary Ric complex are required for the formation of the stable RicT-Rny complex. We demonstrate that proteins of the degradosome-like network of *B. subtilis*, which also interact with Rny, are dispensable for processing of the *gapA* operon. Thus, Rny participates in distinct RNA-related processes, determined by its binding partners, and a RicT-Rny complex is likely the functional entity for *gapA* mRNA maturation.

**IMPORTANCE:** The action of nucleases on RNA is universal and essential for all forms of life and includes processing steps that lead to the mature and functional forms of certain transcripts. In *B. subtilis* it has been shown that key transcripts for energy producing steps of glycolysis, for nitrogen assimilation and for oxidative phosphorylation, all of them crucial processes of intermediary metabolism, are cleaved at specific locations, resulting in mRNA stabilization. The proteins required for these cleavages in *B. subtilis* (Rny (RNase Y), RicA (YmcA), RicF (YlbF) and RicT (YaaT)) are broadly conserved among the firmicutes, including in several important pathogens, hinting that regulatory mechanisms they control may also be conserved. Several aspects of these regulatory events have been explored: phenotypes associated with the absence of these proteins have been described, the impact of these absences on the transcriptome has been documented, and there has been significant exploration of the biochemistry and structural biology of Rny and the Ric proteins. The present study further advances our understanding of the association of Ric proteins and Rny and shows that a complex of Rny with RicT is probably the entity that carries out mRNA maturation.

---

In bacteria, most mRNA species are rapidly degraded, enabling prompt adjustment of translation to changing circumstances. In addition to degradation, some transcripts are matured by specific cleavage events (1). These include the processing of tRNA and ribosomal RNA precursors (2) and selective cleavages that stabilize some transcripts for key metabolic pathways. In *Bacillus subtilis* and related firmicutes, the *gapA* operon encodes five enzymes for triose metabolizing, energy-yielding steps of glycolysis. This operon is regulated transcriptionally by the CggR repressor and post transcriptionally by a cleavage event near the 3’ end of the *cggR* coding sequence (Fig. 1A) (3-6). Cleavage has two consequences; it prevents the synthesis of active CggR, thus derepressing transcription of the operon, and it stabilizes the mRNA sequences that encode downstream genes. These combined effects increase production of glyceraldehyde-3-phosphate dehydrogenase, encoded by the downstream *gapA* gene (4), and presumably also of other enzymes for the glycolytic metabolism of triose phosphates, also encoded downstream. This *gapA* mRNA maturation theme is repeated with the *glnRA* and *atpIBE* transcripts; specific cleavage events stabilize the transcripts (6). In addition to the thematic resemblance of these processing events, which affect central carbon and nitrogen metabolism, as well as oxidative phosphorylation, they are also mechanistically related; all three require Rny and all three require RicA (YmcA), RicF (YlbF) and RicT (YaaT) (6, 7). Rny is a membrane-localized RNase, which is responsible for the initiation of bulk mRNA turnover in *B. subtilis* and many other firmicutes, playing a role analogous to that of RNase E in *E. coli* (8). Both Rny and RNase E are membrane proteins but exhibit little sequence similarity and differ in their domain organizations. Even their membrane anchoring differs; Rny has an N-terminal transmembrane helix (7, 9) and RNase E is attached by an amphipathic helix (10).

**Fig. 1.**
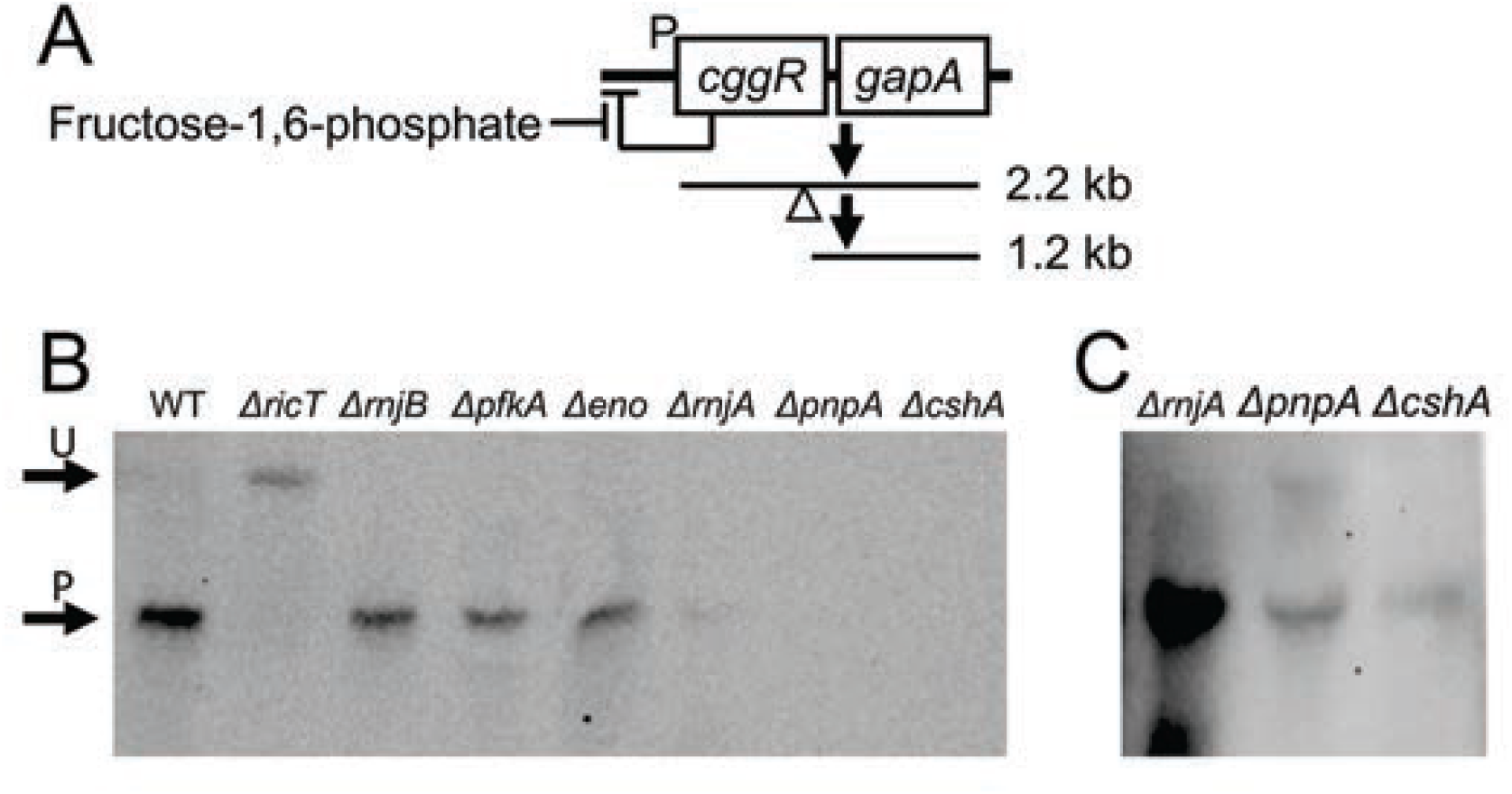
The DLN proteins are not needed for the processing of *gapA* mRNA. Panel A is a diagram explaining the transcripts that are visible in the Northern blot of panel B. The large read through transcript that includes coding sequences for *pgk, tpi, pgm* and *eno* is not depicted for simplicity. The empty triangle shows the approximate position of the cleavage site within *cggR*. Panel B shows the positions of the unprocessed 2.2 kb *cggR gapA* transcript (U) and the 1.2 kb product encoding *gapA* (P) in a Northern blot. RNA preps from wild-type, *ΔricT, ΔrnjB, Δpfk, Δeno, ΔrnjA, ΔpnpA* and *ΔcshA* strains were run on the gel and probed with *gapA* RNA. The image in panel C, shows an over-exposed image of the *ΔrnjA, ΔpnpA* and *ΔcshA* lanes from the same gel.

The three Ric proteins were originally identified as needed for sporulation, competence, and the formation of biofilms in *B. subtilis*; (11-13). Genetic and biochemical evidence suggested that these proteins are needed for the phosphorylation of the master regulator Spo0A, thereby contributing to these developmental pathways (14-16). This, however, is a moonlighting function; *B. subtilis* has hijacked these proteins for the regulation of Spo0A-P formation. In fact, the Ric proteins are present in firmicutes that lack Spo0A (15, 17), and so the more phylogenetically general role for the Ric proteins appears to be in RNA metabolism, particularly for the maturation and stabilization of the above-mentioned transcripts encoding metabolic enzymes, but also for the processing or degradation of several non-coding RNAs (6, 17). RicA, F and T form a 1:1:1 ternary complex and the structure of a RicA:RicF dimer has been solved (14, 15). The ternary complex carries two [4Fe-4S]^+2^ clusters; cluster 1 is coordinated by four cysteine residues on RicT and cluster 2 by cysteine residues contributed by each of the three protomers. The clusters are essential for maturation of the *gapA* transcript (18).

Based on pull-down experiments followed by mass spectrometry it appeared that RicF and RicT are associated with Rny, although in both cases single samples were tested only once (14, 17). Deloughery et al (17) detected RicF but not RicT or RicA in association with Rny, while Carabetta et al (14) detected Rny in a pull-down with RicT, but not with pulldown of RicF or RicA. Based on these data, and on bacterial two hybrid experiments showing the association of RicA and RicF with Rny, it was proposed that the entire Ric complex associates with Rny (6). Because of the inconsistency in these results, it was important to revisit the association of Ric proteins with Rny.

In *Escherichia coli* a multi-component degradosome complex is involved in the global turnover of mRNA, with RNase E acting as a scaffold for assembly of the complex (19). In *B. subtilis* and other firmicutes, binary interactions of several proteins with Rny have been detected (19-21), some of them orthologs of the *E. coli* proteins that interact with RNase E. In *B. subtilis*, these interactions have generally been detected after *in vivo* cross-linking and are proposed to be transient or low affinity in nature, leading to the useful concept of a Degradosome-Like Network (DLN) (22). Despite the transient nature of these interactions, the DLN proteins other than Rny are candidates for involvement in *gapA* maturation. To date, only polynucleotide phosphorylase among the DLN proteins has been tested and found to be dispensable for maturation (23).

Here we have re-investigated the association of Rny with the Ric proteins and have found that only RicT forms a stable complex with Rny, while all three Ric proteins are required to form this stable association. Furthermore, the two Fe-S clusters are needed for the formation of the stable RicT-Rny complex. Based on these data we propose that the ternary complex delivers RicT to Rny, which is likely to be the functional entity for the maturation of the specific mRNAs described above. Here we also demonstrate that the DLN proteins other than Rny are not needed for processing of the *cggR gapA* transcript. Thus, it appears that the roles of Rny in *gapA* maturation and mRNA turnover are distinct; in the first role it associates with RicT and in the second with proteins that comprise the DLN.

## RESULTS

### Among the degradosome proteins only Rny is needed for *gapA* maturation

To understand the mechanism of processing, it is important to identify the proteins involved and clearly the DLN components, which interact with Rny, comprise a set of candidates. It had been shown that a *Δrny* strain failed to process *cggR gapA* or *glnRA* transcripts (6, 20) leading to the plausible conclusion that Rny is the nuclease that accomplishes processing. But the endoribonucleases RnjA and RnjB, and the RNA helicase CshA, as well as Eno and Pfk remained as possible candidates for involvement in stabilizing cleavage events (SCEs). To determine if they are needed for processing, we used a set of deletion mutants with inactivated DLN genes, all of which were verified by sequencing. Northern blotting was carried out using a probe complementary to the coding strand of *gapA*. Fig. 1B demonstrates that in the absence of enolase, phosphofructokinase, RNase J1 or RNase J2, maturation took place; the bulk of the *gapA* signal is at the position of processed product in the wild-type strain. In contrast, the signal from a *ricT* deletant control is unprocessed, as expected. In repeated experiments we have noted that in the *ΔrnjA, ΔcshA*, and *ΔpnpA* strains the total *gapA* signal was extremely low, not surprising given the pleiotropic effects of these mutations (26-28). After prolonged exposure of the Northern blot, a signal corresponding to processed *gapA* mRNA was detected in the *ΔrnjA, ΔpnpA* and *ΔcshA* lanes (Fig. 1C). The dispensability of PnpA for this maturation event was reported previously (23). We conclude that DLN proteins other than Rny are not absolutely required for *cggR gapA* processing, focusing attention on the Ric proteins as we explore the mechanism of Ric-dependent mRNA processing.

### Strategy and validation of the approach

To clarify the interactions of Rny with Ric proteins, we carried out pull-down experiments in which one protein was tagged C-terminally with a 3xFLAG (3FL) epitope and anti-FLAG magnetic beads were used to pull down associated proteins. The strains expressing the FLAG-tagged Ric proteins used in all the experiments described below were in backgrounds deleted for the cognate wild-type genes and are thus the only source of these proteins. Rny-3FL was inserted at the native locus, under its normal control, while the three Ric-3FL constructs were placed in the ectopic *thrC* locus under control of the inducible P*spac* promoter. Fig. S1 verifies that the Ric proteins are induced by the addition of isopropyl β-D-1-thiogalactopyranoside (IPTG). Importantly, Fig. S2 presents a Northern blot verifying that the tagged Ric and Rny proteins support processing of the *gapA* transcript. For the pull-down experiment, bound proteins were detected by Western blotting using either anti-Rny (a kind gift from J. Stulke) or anti-Ric antisera, which was raised in Rabbits against the purified ternary Ric complex and detects all three Ric proteins (18). Verification of the anti-Ric antiserum with individual *ric* deletion strains are shown in Fig. S3 A. Fig. S3B likewise shows that the anti-FLAG and anti-Rny antisera correctly identify Rny-3FL and Rny. Fig. S3C demonstrates the ability of the beads to pull down Rny-3FL, incidentally verifying that the inclusion of Triton X100 has solubilized this tagged membrane protein. The lysates for these experiments were prepared without cross-linking and in the presence of added RNase to minimize possible bridging by bound RNA.

### Rny stably interacts with RicT but not with RicA or RicF

To investigate whether the Ric ternary complex associates with Rny, perhaps to carry out mRNA maturation (6), we first used a strain expressing Rny-3FL as the only source of Rny, together with the control wild-type strain. These strains were grown to late log phase in LB broth, and lysates were incubated with anti-FLAG magnetic beads after solubilization of membrane proteins, including Rny, by the addition of Triton X100 (1%, w/v) as verified by Fig. S3C. After washing, bound proteins were eluted using an excess of FLAG peptide. Fig. 2A shows that RicT, but not RicA or RicF were pulled down with Rny, although in some experiments, traces of RicF and RicA were visible. The load lane showed that RicA and RicF were abundantly present (Fig. 2A, lane L) and the absence of these two proteins in the 18-fold concentrated eluants is therefore highly significant, particularly because the anti-Ric antiserum reveals a stronger signal for RicA and RicF than for RicT, as is evident in Fig. 2A lane L. The presence of RicT in the eluant (lane E) and its absence in the last wash lane (LW) shows that the association of RicT with Rny was stable during this procedure. It appears then, that Rny exists in a stable complex with RicT, but that RicA and RicF are absent from this complex. Fig. S3D and E show that when an identical pull-down experiment was performed with a wild-type lysate, lacking tagged Rny, neither the Ric proteins nor Rny are recovered in the eluant fraction, showing that these proteins do not bind significantly to the anti-FLAG beads.

**Fig. 2.**
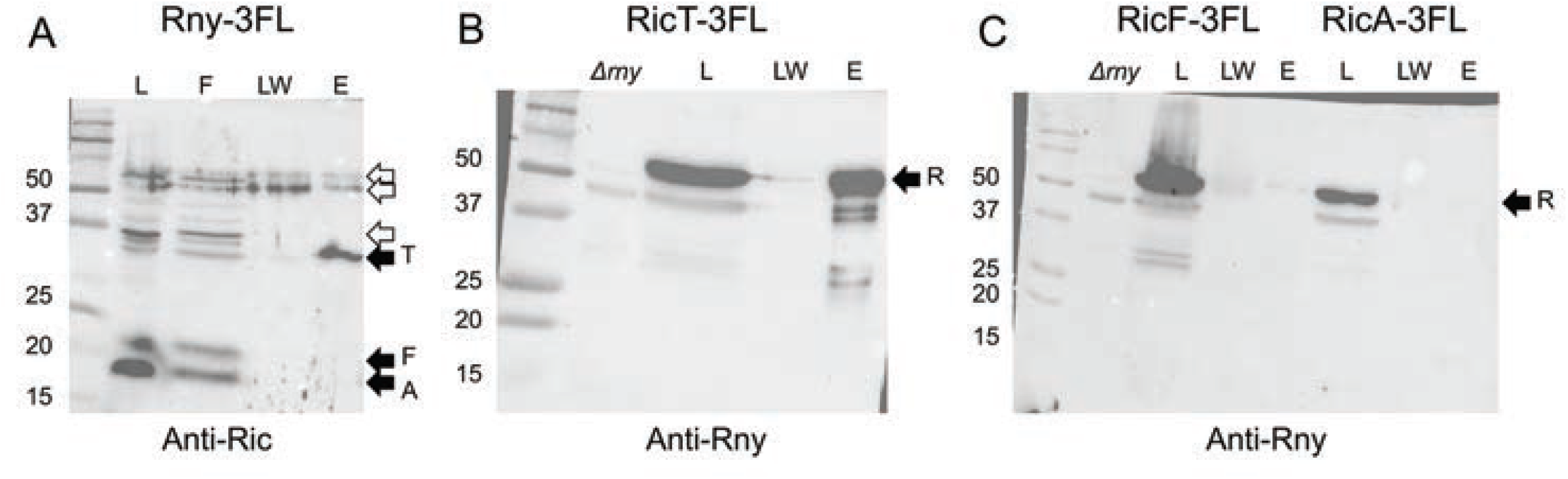
RicT and Rny pull one another down. In panel A an Rny-3FL lysate was incubated with anti-FLAG magnetic beads and the load, flow through, last wash and eluant fractions were analyzed by Western blotting with anti-Ric antiserum. In panel B, a RicT-3FL lysate was similarly treated and the load, last wash and eluant fractions were blotted with anti-Rny antiserum. Panel C documents the failure of RicF-3FL and RicA-3FL to pulldown Rny. In panels B and C a *Δrny* lysate is shown to verify the position of the Rny signal and its presence in the lysate. In the three panels, the solid arrows show the positions of RicT, RicF, RicA and Rny, while the empty arrows show prominent cross-reacting bands. The eluant samples were about 18-fold concentrated compared to the loads.

The eluant samples from this experiment were analyzed by mass spectrometry, which confirmed the presence of RicT but showed no significant presence of RicA or of RicF (Table S1). Notably, none of the degradosome network proteins were present significantly above background, although a number (CshA and RnjB) showed marginally more presence in the Rny-3FL than in the wild-type pull-downs. These results do not conflict with the many reports that PnpA, RnjA, RnjB, CshA, Eno and PfkA contact Rny, but rather argue that the various DLN interactions are not as stable as the RicT interaction with Rny, under the present conditions.

To verify the association of Rny with RicT, we used a 3xFLAG-tagged RicT (RicT-3FL) construct in a strain carrying a *ΔricT* mutation in the native locus. As shown in Fig. 2B, RicT-3FL pulls down Rny, detected using anti-Rny antiserum. Fig. S4 shows that as expected, RicT-3FL also pulls down RicA and RicF, showing that it is competent for the formation of the ternary complex. Therefore, RicT exists in two stable complexes; in association with Rny and with RicA and RicF, as had been reported previously. In fact, Fig. S4 also shows that each tagged Ric protein pulls down the other two, showing that the Ric complex is formed normally with all three C-terminally tagged proteins, as noted before with a YFP tag (14). In a further confirmation of these results, we used FLAG-tagged RicA (RicA-3FL) and RicF (RicF-3FL). Although RicA-3FL and RicF-3FL pulled down RicT, and respectively pulled down RicF or RicA (Fig. S4), no Rny was detectable in the eluants (Fig. 2C). Taken together, these experiments demonstrate that RicT and Rny are stably associated with one another, and that RicA and RicF are in a complex with RicT but are not detectably associated with Rny. We obtained no evidence for a stable association of Rny with the ternary RicAFT complex.

### RicA and RicF facilitate the association of RicT with Rny

What then are the roles of RicA and RicF, which are required for mRNA processing? Because RicA and RicF are required for maturation, and are present in a ternary complex with RicT, we asked if they are needed to form the Rny-RicT association. As shown in Fig. 3, elimination of either of these proteins prevented the pull-down of RicT by Rny-3FL. We conclude that RicA and RicF function *in vivo* to aid in the formation of an Rny-RicT binary complex.

**Fig. 3.**
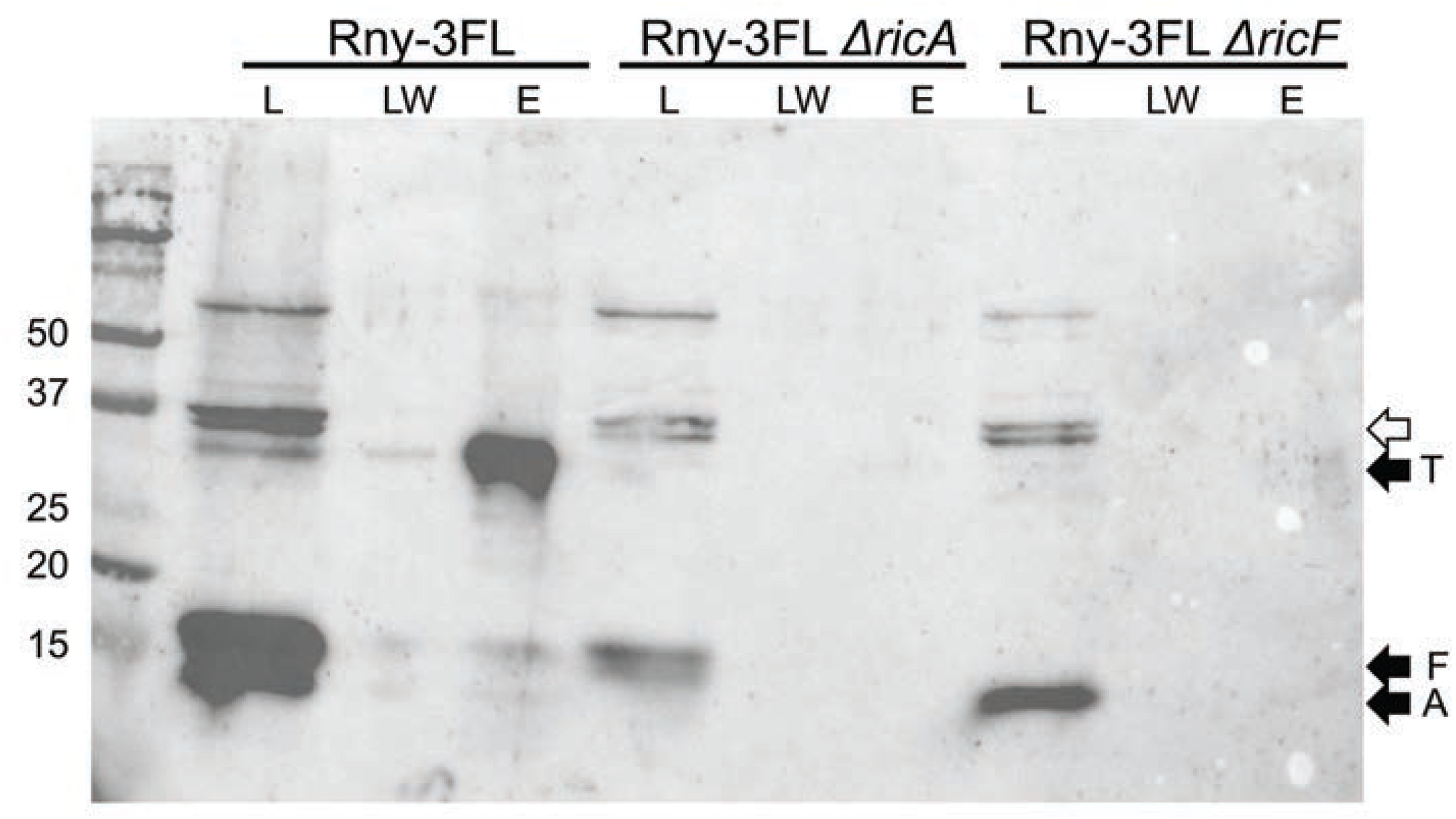
RicA and RicF assist in the formation of the Rny-RicT complex. From the Rny-3FL lysate, RicT is evident in the eluant (E) lane. In Rny-3FL *ΔricA* and Rny-3FL *ΔricF* backgrounds, RicT is not detectable. Load (L), last wash (LW) and eluant (E) samples are shown for each lysate. The blot was developed with anti-Ric antiserum. The empty arrow shows the position of prominent cross-reacting bands. The eluant samples were about 18-fold concentrated compared to the loads.

### Roles of the Fe-S clusters in the formation and stability of the Rny-RicT complex

As noted above, the ternary Ric complex carries two [4Fe-4S]^+2^ clusters. Cluster 1 is entirely coordinated by cysteine residues on RicT, while cluster 2 is ligated to residues donated by each of the proteins. We have shown that both clusters are required for *gapA* maturation (18) and we now asked if they are needed for RicT to associate with Rny. As shown in Fig. 4A, the pull-down of RicT by Rny-3FL is markedly decreased by both the C161S and C198S mutations of RicT that prevent formation of cluster 1 and the C167S mutation of RicT that prevents formation of cluster 2. These results suggest that the clusters are required for either the formation or the stability of the RicT-Rny complex once formed, or for both. In this figure and in Fig. 3, traces of RicA and F are present in the wild-type eluant fraction, which is highly concentrated compared to the load. This may be non-specific background or it may represent a transient form involving all four proteins, as discussed below.

**Fig. 4.**
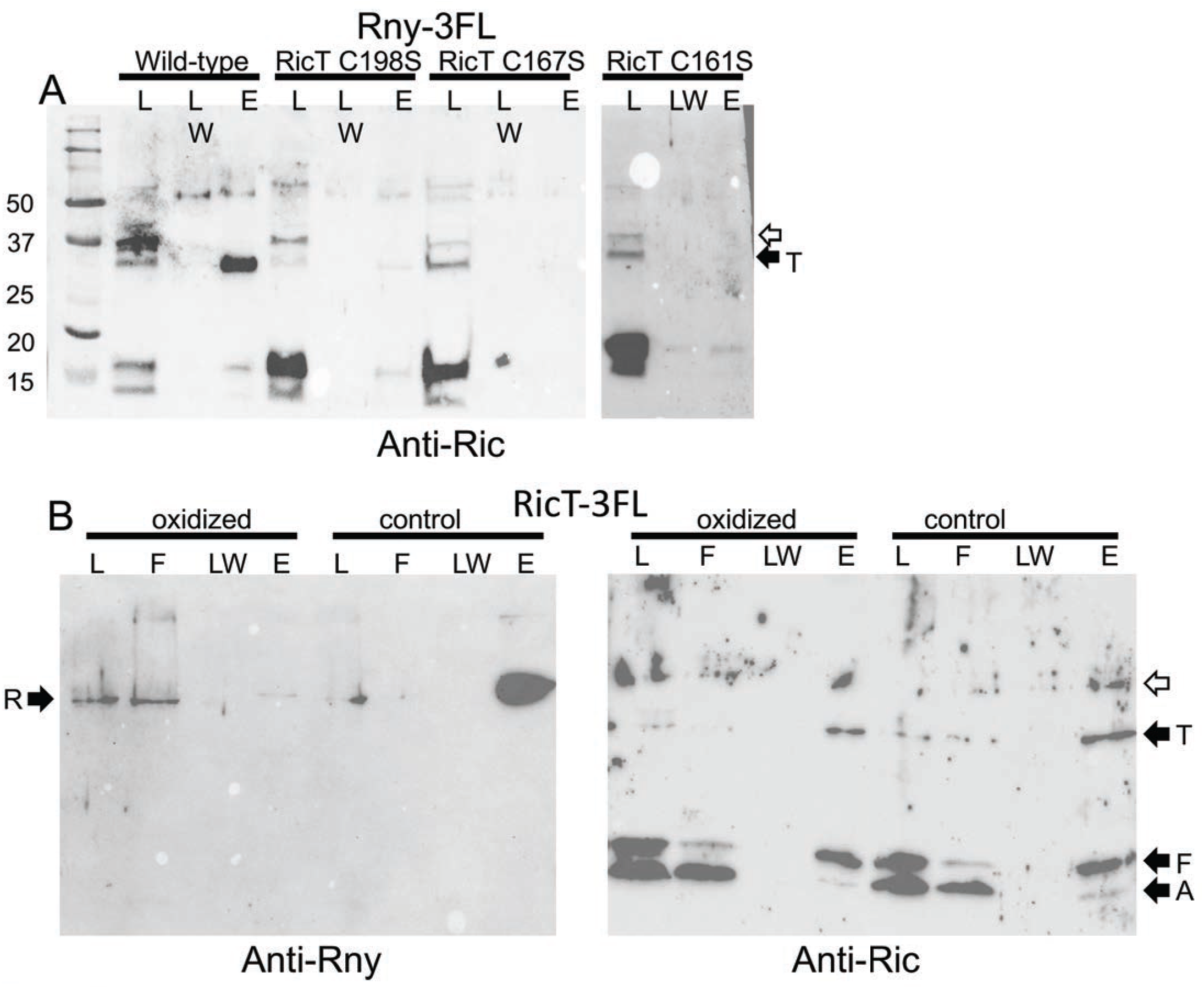
The iron sulfur clusters are needed for formation and stability of the Rny-RicT complex. (A) Rny-3FL strains expressing wild-type RicT, RicT C198S, RicT C167S or RicT C161S mutant proteins were used for pull-downs and the blots were developed with anti-Ric antiserum. The first two mutations affect cluster 1 and the C161S mutation affects cluster 2. The two images in this panel are from different gels. Panel B shows that oxidation and chelation treatment of a lysate prevents pulldown of Rny by RicT-3FL, shown with anti-Rny antiserum, but not pulldown of RicA and RicF, shown with anti-Ric antiserum. The empty arrows indicate prominent cross-reacting bands.

To determine if the RicT cluster 1 is needed for the stability of RicT association with Rny *after* the complex is formed, we carried out a pull-down experiment using RicT-3FL following a treatment expected to destroy the cluster. As reported previously, both oxygen-sensitive Fe-S clusters can be completely stripped from the ternary Ric complex by oxidation and chelation using potassium ferricyanide plus EDTA, without compromising the stability of the complex (15). For this experiment, lysates of a RicT-3FL strain were divided into two portions, and then incubated with and without the addition of EDTA and potassium ferricyanide before proceeding with the usual pull-down protocol. As shown in Fig. 4B, this treatment nearly completely prevented the pull-down of Rny, suggesting that although clusters 1 and 2 are needed for formation of the Rny-RicT complex, cluster 1, which is entirely coordinated by RicT residues, is needed to maintain the complex. In contrast, the pull-down of RicA and RicF by RicT-3FL was not affected by oxidation and chelation (Fig. 4B), consistent with previous results showing that the ternary complex did not need the clusters for stability (15, 18). Although we do not have direct confirmation that chelation plus oxidation has destroyed the clusters in the Rny-RicT complex, this seems highly probable because the identical procedure strips the clusters from the purified Ric ternary complex *in vitro*. Thus, at least cluster 2 and most likely both clusters are needed for formation of the Rny-RicT complex and cluster 1 is likely needed for stabilization of the complex, once formed.

## DISCUSSION

This study shows that RicA and RicF deliver RicT to form a stable complex with Rny. We propose that this takes place during a transient interaction of the ternary complex with Rny, forming Rny-RicT, most likely the functional entity for RNA processing. Although we have not presented direct experimental evidence that the Rny-RicT complex is competent for RNA processing, this model explains why all three Ric proteins are needed for processing even though RicA and RicF do not associate stably with Rny, the relevant ribonuclease. Likewise, the finding that transfer to Rny requires the Fe-S clusters is consistent with the essentiality of the clusters for processing (18). The conclusion that RicT-Rny is the active form for RNA maturation is strongly supported by the nearly universal presence of RicT among the firmicutes, in contrast to RicA and RicF, which are more limited in their distributions (15). Finally, the idea that RicT-Rny is the functional entity is consistent with the dispensability of the DLN proteins for *gapA* transcript processing; these are the only additional proteins known to interact with Rny.

The results presented here are in accord with the finding by DeLoughery et al, showing that RicT is normally associated with the cell membrane, but becomes cytosolic when Rny or RicF are absent. RicT is about 10-fold more abundant *in vivo* than RicA and RicF and is soluble *in vitro* as a monomer (18). The RicF-RicA heterodimer is also soluble and thus there may be three forms of RicT; monomer, in the ternary complex, and in association with Rny. There may also be two forms of RicA-RicF; the heterodimer and as part of the ternary complex. RicA and RicF may in fact be viewed as shuttle proteins, cycling RicT from a pool of monomers to Rny via the ternary complex.

In this study, we have not demonstrated direct transfer of RicT from RicAFT to Rny, although this is the most likely mechanism. Direct transfer of RicT from the ternary complex to Rny is supported by the requirement of cluster 2 for formation of the RicT-Rny complex, because this cluster is coordinated by all three Ric proteins. Direct transfer of RicT from the ternary complex would require the existence of a short-lived quaternary intermediate before the departure of RicA-RicF. This may explain an apparent contradiction of our results with the literature. Deloughery et al have reported an association of Rny with RicA and RicF using a bacterial two hybrid system (17). This method may report transient interactions because it depends on brief bursts of transcription, which then result in the accumulation of a stable, detectable product. Indeed, in some experiments we have noted the presence of small amounts of RicA and RicF in the eluant fractions when Rny-3FL is pulled down (see for example Figs. 3 and 4), consistent with such a transient intermediate. It will be important to verify the direct transfer mechanism using purified proteins.

Just as RicT exists in at least two forms in the cell; in association with RicA and RicF and bound to Rny, Rny also exists in multiple forms; bound to RicT, as shown here, and transiently associated with other members of the degradosome network. The DLN interactions have been detected in some cases after cross-linking, over-expression of one of the proteins involved or with purified proteins *in vitro* (19-21, 29). However, we have not detected them by LC-MS-MS in our pull-down using Rny-3FL (Table S1). Our data are thus consistent with the emerging picture of the DLN as a dynamic network of binding partners, characterized by weak interactions. We have repeated this analysis without the addition of RNase to the lysates to determine if the absence of such signals in our experiment was because we had removed bridging RNA species, but the same result was obtained. The picture of a DLN typified by transient Rny interactions stands in contrast to the apparently stable interaction of Rny with RicT, which is persistent in lysates without cross-linking or over expression. This difference fits well with the non-essentiality of degradosome network components, other than Rny, for the maturation of the *cggR gapA* transcript and clearly distinguishes the maturation and global turnover roles of Rny. We propose then, that Rny-RicT is the functional RNase complex for *cggR gapA* maturation and probably more generally, for *glnRA* and *atpIBE* as well.

Much remains unknown. On important issue concerns the role of RicT. Rny appears to have an inherent preference for single stranded RNA, preferably cutting after a G residue (30, 31), and the role of RicT may be to direct Rny to certain transcripts, rather than to specific sites. Of particular interest for further studies are the roles of the Fe-S clusters. Although the clusters are not needed to hold the ternary complex together (15, 18), they may still play structural roles, such as stabilizing a loop within RicT (cluster 1) and the C-terminal flexible segments of RicF and RicA (cluster 2) (18). And, the clusters may also be involved in regulatory signaling.

*cggR gapA, atpIBE* and *glnRA* mRNA processing events presumably enable efficient glycolysis, oxidative phosphorylation and the synthesis of glutamine. Glutamine synthetase (GlnA) is a so-called trigger enzyme (32), playing a crucial regulatory role in the actions of GlnR and TnrA, the latter a regulator of global nitrogen metabolism (33). Thus, these processing events are poised at strategic control points for the life of *B. subtilis* and its relatives.

## MATERIALS AND METHODS

### Plasmids and strains

All the *B. subtilis* strains are in the IS75 (*his leu met*) background, a derivative of the domesticated laboratory strain 168. Strains were constructed using transformation and are listed in Table S2. Oligonucleotide primers are listed in Table S3.The Rny-3FL construct was made by cloning the 3’ end of *rny* into the *Eco*RI and *Bam*HI sites of pGP1331 (19), which places the Rny sequence in-frame with 3xFLAG. The primers used for producing this fragment by PCR were Rny-F and Rny-R. The resulting plasmid was then inserted in the *rny* locus by single reciprocal recombination. The three Ric-3FL constructs were inserted in the ectopic *thrC* locus under control of the IPTG-inducible P*spac* promoter. In step one, pGP1087 (34) was opened with *Kpn*I and *Cla*I, and then *ricT*, produced by PCR with the primers RicT-F1 and RicT-R1, was inserted into the opened vector using Gibson Assembly. The resulting plasmid, pGP1087-RicT, contained the P*spac* promoter in front of *ricT* fused to 3xFLAG with *lacI* downstream. A set of Gibson Assembly (New England Biolabs, Inc.) primer pairs (RicT-F2 and RicT-R2) was then used to isolate this *ricT* fragment and insert it between the *Bam*HI and *Hin*dIII sites of pDG1664 (35) to produce pDG1664-RicT. pDG1664 is an integrative plasmid for insertion into *thrC*. For construction of the comparable IPTG-inducible *ricA* and *ricF* integrative plasmid, pDG1664-RicT was opened with *Kpn*I and *Cla*I and the primer pairs RicA-F+RicA-R and RicF-F+RicF-R were used to insert *ricF* and *ricA* in-frame with 3xFLAG after cutting the PCR products with *Kpn*I and *Cla*I. The resulting plasmids were used to insert the constructs into the chromosomal *thrC* locus by replacement recombination with selection for erythromycin-resistance. *RicA* was placed under IPTG control together with the upstream *ymcB* gene because the two open-reading frames appear to be translationally coupled.

### Pull-down experiments

Strains for pull-down experiments were grown in Luria-Bertani broth containing 1 mM IPTG, to late log phase. Cells were harvested by centrifugation and washed twice in lysis buffer (5 ml of HEPES buffer (20 mM pH 7.4), 0.2 M NaCl, 1 mM MgCl_2_, 2 μM CaCl_2_). The final pellets were frozen and held at -80 C until use. The pellets were resuspended in 2 ml of the same buffer to which was added DNase (10 ug/ml), RNase (10 ug/ml), lysozyme (200 ug/ml) and 1 tablet per 5 ml of buffer of EDTA-free protease inhibitor (Pierce). The suspensions were incubated at 37 C for 15 minutes and 5 mM PMSF was added. The suspensions were then chilled in 15 ml Falcon tubes and sonicated 3 x for 30 seconds on ice and allowed to cool between treatments for 1 minute. 0.2 ml of 10% Triton X100 was added to each lysate to reach a final concentration of 1% (w/v). The lysates were then centrifuged at 6000 RPM to remove unbroken cells and some debris. The pull-downs were carried out immediately on these supernatants. A portion of the lysate was retained as the load fraction. Anti-FLAG functionalized magnetic beads (Thermofisher A36797) were brought to room temperature and gently mixed to homogeneity. To an Eppendorf tube, 200 μl of the bead slurry was added and 1 ml of 20 mM HEPES, pH 7.4, 200 mM NaCl, Triton (0.2%) was added. After mixing, the suspension of beads was placed on a magnetic rack and the supernatants removed. This wash step was repeated for a total of 2 washes. To the washed beads, 1.8 ml of clarified lysate was added and the slurry was mixed gently and then incubated for 30 minutes at room temperature with gentle shaking. The slurry was again placed on the magnetic rack and the supernatant retained (flow through fraction). 1.0 ml of PBS was added to the tube and after mixing, the samples were placed on the magnet and the supernatants were discarded. This was repeated 3-times and the final supernatant was retained as the final wash sample. For elution from the beads, 100 μl of FLAG peptide (Thermofisher A36805) dissolved in PBS to a final concentration of 1.5 mg/ml, was added. The samples were mixed and incubated at room temperature for 10 minutes with intermittent mixing. The supernatant was removed on the magnetic rack and retained as the eluant fraction.

### SDS-Polyacrylamide and Western blotting

SDS-PAGE and Western blotting were carried out as described (18), except that conventional 12% Laemmli gels were used instead of Bis-Tris gel buffer. After electrophoresis was performed, the gels were blotted to nitrocellulose membranes using a Transblot Turbo apparatus (Bio-Rad) and developed using rabbit antisera raised against the ternary Ric complex, commercial anti-FLAG M2 monoclonal antiserum (Sigma) or a rabbit antiserum raised against Rny, a gift from Jörg Stulke. The signals were visualized using ECL Prime Western blotting detection reagent (GE Healthcare).

### Northern blotting

Northern blotting for the *cggR gapA* transcript was performed as described (18), using a probe prepared by *in vitro* transcription of a sequence complementary to the *gapA* coding strand. Samples were processed in a FastPrep FP120 instrument for 3 cycles (45, 45, and 30 s) at a speed setting of 6.0. The samples were treated with 1 μl of RNase-free DNase (NEB) to 100 μl of RNA solution and cleaned up with a Monarch RNA cleanup kit (NEB). RNA concentrations were measured with a NanoDrop 1000 spectrophotometer (Thermo Scientific). RNA was resolved using agarose gel electrophoresis in 3-(N-morpholino) propanesulfonic acid (MOPS) buffer (0.2 M MOPS, 20 mM sodium acetate, 10 mM EDTA) with 1.5% agarose containing 5% formaldehyde. Electrophoresis was performed at 50 V for 50 min. The RNA loading buffer contained 50 μl 80% glycerol, 200 μl formaldehyde, 240 μl 10x MOPS, and a few crystals of bromophenol blue. All solutions were made with diethyl pyrocarbonate-treated water. For Northern analysis, ethidium bromide was omitted. RNA was transferred to Bright Star-Plus positively charged nylon membranes (Ambion) using passive (capillary) transfer. The membrane was subjected to UV cross-linking in a Spectrolinker XL-1000 UV cross-linker. The membrane was then prehybridized with DIG Easy Hyb (Sigma-Aldrich), hybridized with digoxigenin (DIG)-labeled RNA probe, washed, and incubated with anti-digoxigenin-AP FAB fragments (Sigma-Aldrich), and the bands were developed using disodium 3-[4-methoxyspiro [1,2-dioxetane-3,2’-(5’-chloro)tricyclo (3.3.1.13,7) decan]-4-yl]phenyl phosphate (CSPD; Sigma-Aldrich}, all per the manufacturer’s instructions. The digoxigenin-labeled RNA probes were transcribed from PCR products containing a T7 RNA polymerase promoter fused to *gapA* sequences with T7 RNA polymerase (New England Biolabs) and DigRNA labeling mix (Sigma-Aldrich) per the manufacturer’s instructions. Importantly, to obtain product it was necessary to supplement the New England Biolabs buffer with 5 mM DTT. The sequences of the *gapA* forward and reverse primers were 5’-CGATGCGCTAACCACGATGTTA and 5’-GAAAT<u>TAATACGACTCACTATAGG</u>GACCATAACCATTGTAGAAAG, respectively. The T7 RNA polymerase promoter sequences are underlined. The bands were visualized and digitized using a Bio-Rad ChemiDoc MP imaging system.

### Treatment with potassium ferricyanide and EDTA

To a lysate prepared as described above from the RicT-3FL strain, EDTA (pH 7.5) was added to 15 mM. The sample was incubated on ice for 5 min. To this, potassium ferricyanide was then added to reach a concentration of 10 mM from a freshly prepared 100 mM stock solution in lysis buffer. Incubation was continued on ice for 1 hour. An identical control sample was incubated similarly with the addition of buffer instead of the two reagents. Pull-downs were carried out as described above.

### Mass spectrometry

Eluant samples from pull-down experiments were run on 12% SDS-PAGE gels. The gels were fixed using methanol (50%) + acetic acid (10%) and stained with Coomassie Simply Blue. After destaining in MillQ water, the eluant lanes were excised and full-length trypsin digestion was performed after dicing the lane into 1 mm cubes. The resulting peptides were analyzed by LC-MS/MS on an Orbitrap Fusion Lumos MS instrument. The MS/MS spectra were searched against the Uniprot *Bacillus subtilis* 168 database using Sequest search engines on the Proteome Discoverer (v2.4) platform. The false discovery rate was less than 1%.

## ACKNOWLEDGEMENTS

We are grateful for discussions and suggestions from all the members of the Dubnau lab as well as from V. Petrou and A. Punetha. This work was supported by NIH Grant RO1GM057720. The mass spectrometry data were obtained from an Orbitrap mass spectrometer funded in part by an NIH Grant NS046593, for the support of the Rutgers Mass Spectrometry Center for Integrative Neuroscience Research.

---

**Fig. S1.**
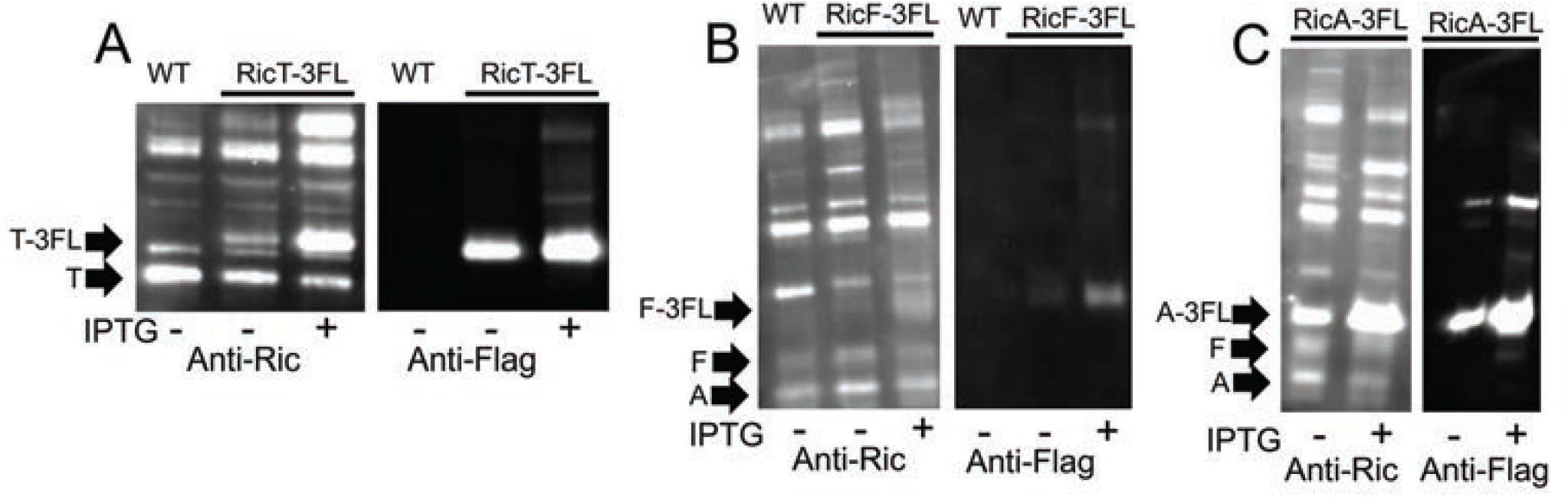
Verification of the Ric-3FL constructs. Panels A, B and C show that the RicT-3FL, RicF-3FL and RicA-3FL express the appropriate fusion proteins, detectable using both anti-Ric and anti-FLAG antisera and that the constructs are IPTG-inducible. These strains did not carry knockouts of the native *ric* genes and thus signals for the untagged proteins were detected.

**Fig. S2.**
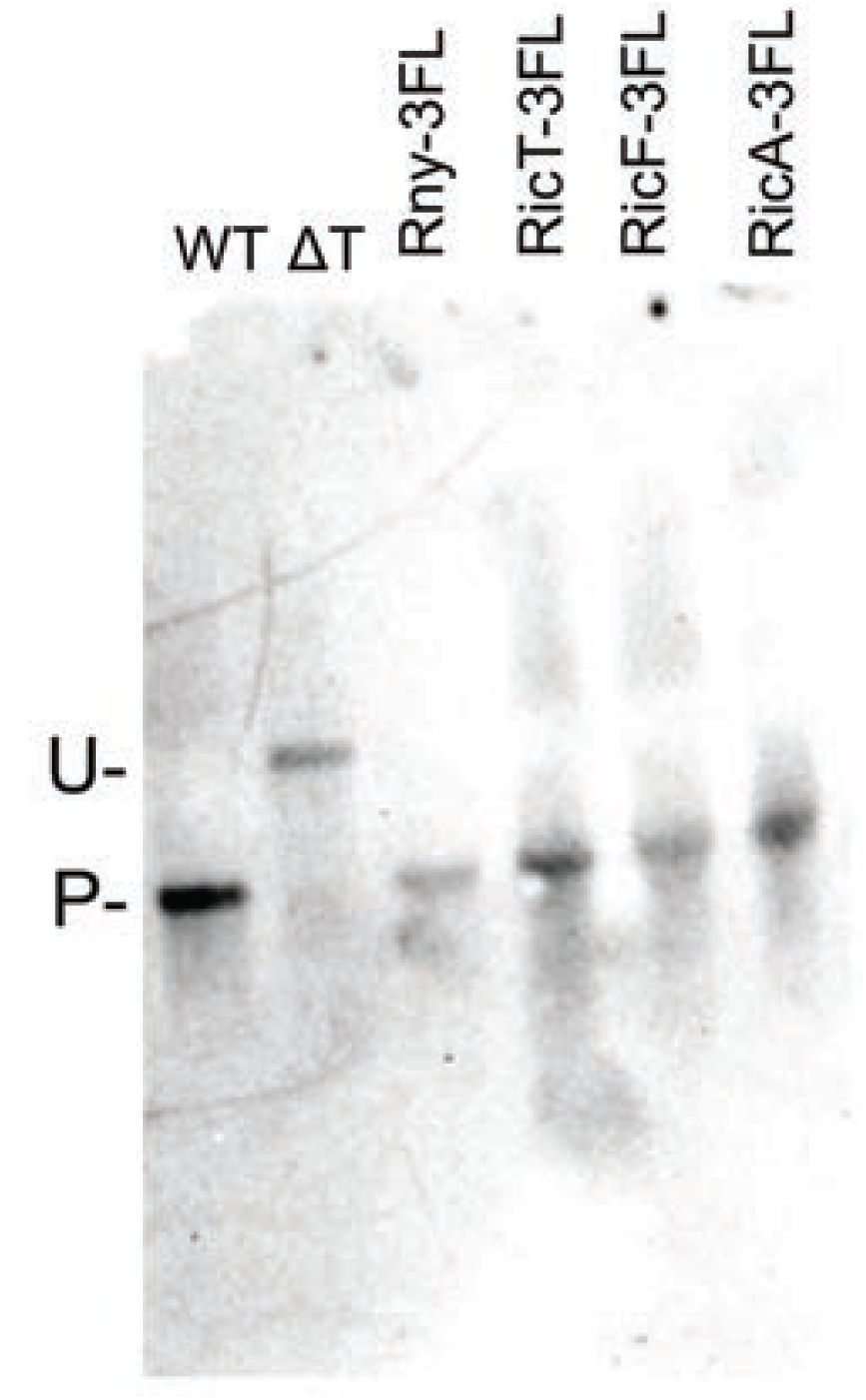
Ric-3FL and Rny-3FL proteins support processing of the *cggR gapA* transcript. Northern blotting was carried out using a probe complementary to the coding strand of *gapA*. Each of the four fusion strains were the only source of RicA, RicF, RicT and Rny in the cells. U and P show the positions of the unprocessed and processed RNA species (∼2.2 and ∼1.2 kb, respectively). The *ricT* deletion and wild-type (IS75) control lanes show the positions of these two species, respectively.

**Fig. S3.**
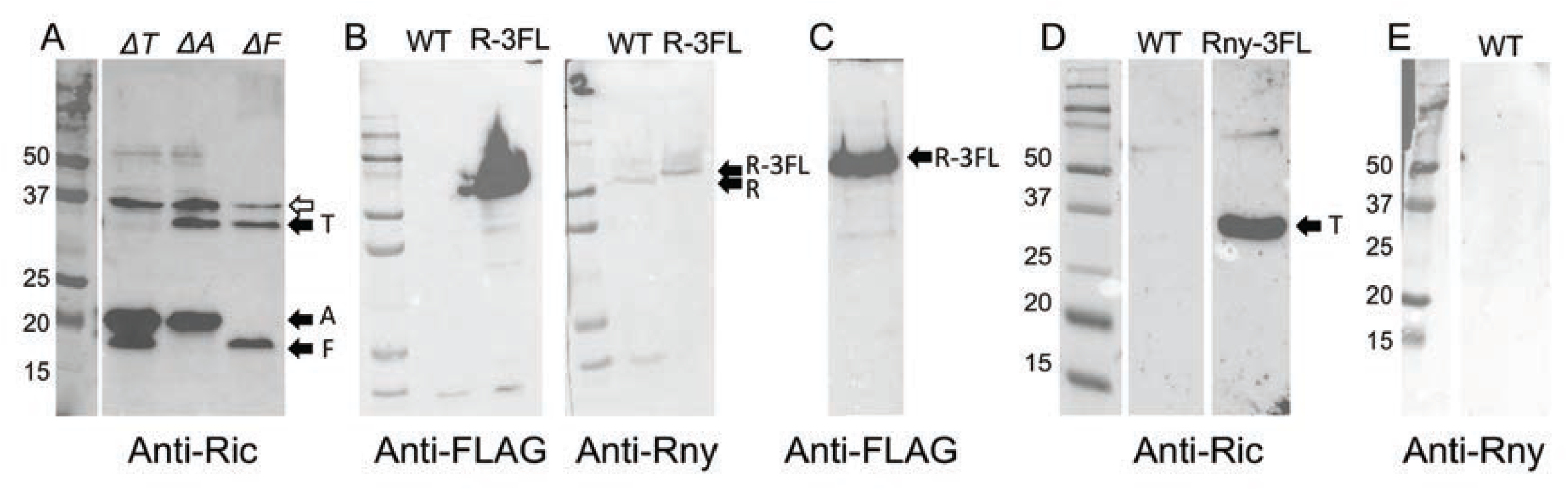
Verification of anti-serum specificities and of the anti-FLAG magnetic beads. (A. The appropriate Ric proteins bands are missing from total lysates of each indicated deletion strain. (B) In a lysate of the Rny-3FL strain, the anti-FLAG and anti-Rny antisera reveal the fusion protein, but only the wild-type Rny signal is evident in a wild-type lysate. (C) An eluant fraction is shown from a pull-down experiment using an Rny-3FL lysate and anti-FLAG antiserum. The magnetic beads successfully pull down Rny-3FL. (D) Eluant fractions from pull-down experiment using wild-type and Rny-3FL lysates. The blots were developed with anti-Ric antiserum. RicT does not bind non-specifically to the beads. (E) No signal is detected in the eluant fraction from a mock pull-down experiment with a wild-type lysate, using anti-Rny antiserum.

**Fig. S4.**
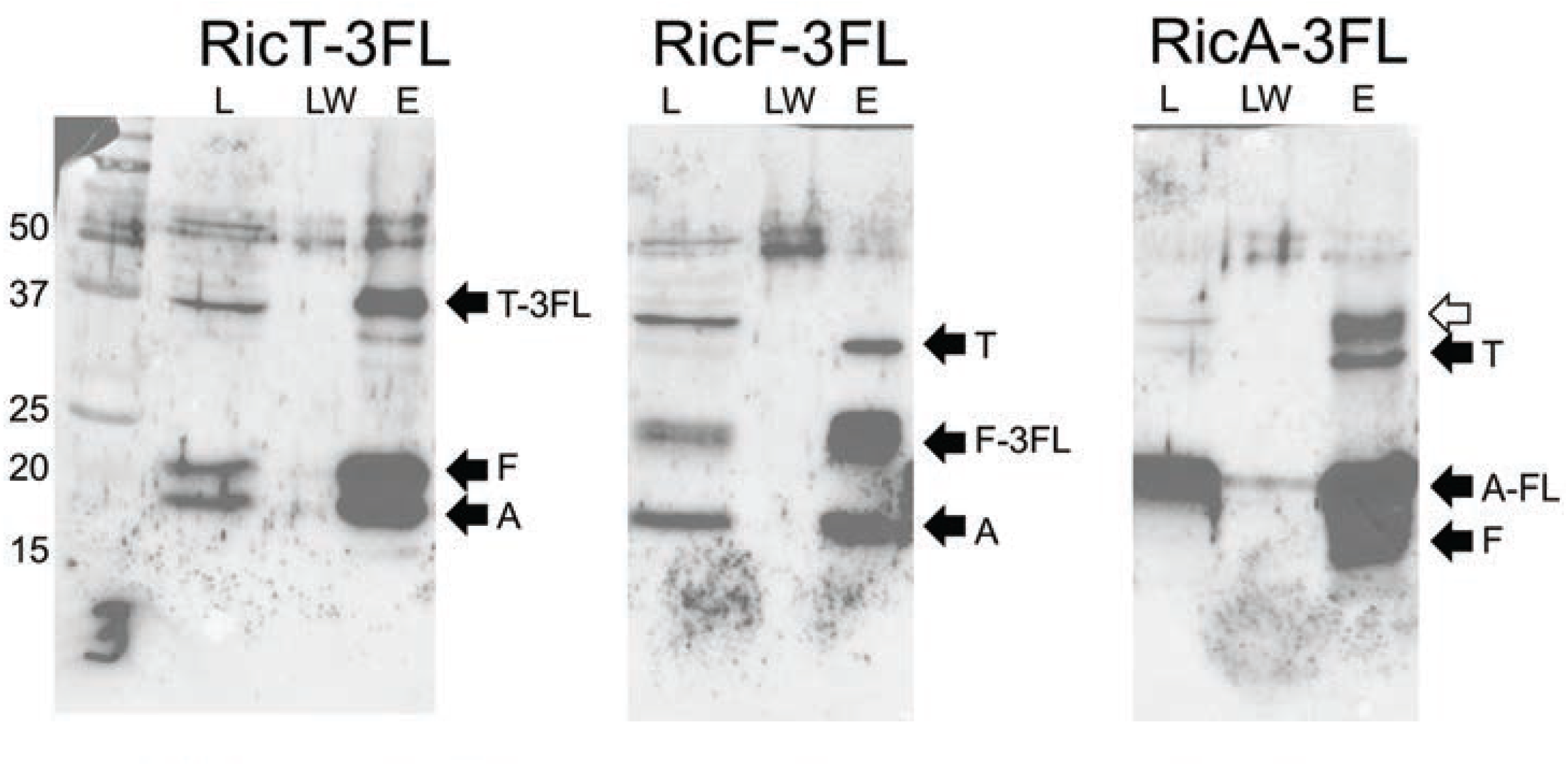
The three Ric-3FL constructs can pull one another down with anti-FLAG beads. In all three panels the blots were developed with anti-Ric antiserum. The empty arrow shows a cross-reacting band.

**Table S1.** Mass Spectrometry Data: Pull-down with Rny-3FL

| Protein | # AA (peptides)<sup>a</sup> | Coverage (%)<sup>b</sup> | Abundance<sup>c</sup> (IS75-1) | Abundance<sup>c</sup> (IS75-2) | Abundance<sup>c</sup> (Rny-3FL-1) | Abundance<sup>c</sup> (Rny-3FL-2) | Ratio<sup>d</sup> | T test |
| --- | --- | --- | --- | --- | --- | --- | --- | --- |
| Rny | 520 (72)) | 85.77 | 2.1 | 3 | 173.5 | 221.4 | 77.2 | 0.01 |
| RicT | 275 (29) | 80.0 | 6.5 | 8 | 191 | 194.5 | 26.6 | 0.00 |
| RicF | 149 (3) | 27.52 | 0 | 0 | 1 peptide<sup>e</sup> | 0 | - | 0.42 |
| RicA | 143 (2) | 18.18 | 49.5 | 114.6 | 94.4 | 141.4 | 1.4 | 0.47 |
| Eno | 430 (28) | 73.26 | 72.1 | 99.4 | 118 | 110.5 | 1.3 | 0.18 |
| PfkA | 319 (31) | 79.62 | 81.5 | 109.9 | 121.7 | 86.9 | 1.1 | 0.74 |
| Pnp | 705 (42) | 56.03 | 82.3 | 105.2 | 114.5 | 98 | 1.1 | 0.47 |
| CshA | 494 (39) | 63.56 | 44.7 | 90.1 | 166.3 | 98.8 | 2.0 | 0.25 |
| RnjB | 555 (44) | 82.34 | 6.4 | 97.1 | 184 | 112.6 | 2.9 | 0.24 |
| RnjA | Not detected |  |  |  |  |  |  |  |

<sup>a</sup>Total number of amino acid residues (predicted number of peptides after trypsinization).<sup>b</sup>Combined identified amino acids/total amino acids in the protein.<sup>c</sup>Reflects the signal strength in the raw data.<sup>d</sup>(Average abundance for Rny-3FL/average abundance for IS75).<sup>e</sup>One peptide was detected in this sample.

**Table S2.** Strains.

Strains
| Strain number | Relevant genotype<sup>a</sup> | Source |
| --- | --- | --- |
| IS75<sup>a</sup> | *his leu met* | Lab strain |
| BD9000 | *rny-3FL (spc)* | This work |
| BD9170 | *P<sub>spac</sub>-ricT-3FL::thr (erm) ricT::kan<sup>b</sup>* | This work |
| BD9171 | *P<sub>spac</sub>-ricF-3FL::thr (erm) ricF::kan<sup>b</sup>* | This work |
| BD9172 | *P<sub>spac</sub>-ricA-3FL::thr (erm) ric::kan<sup>b</sup>* | This work |
| BD9195 | *rny-3FL (spc) ΔricA<sup>c</sup>* | This work |
| BD9196 | *rny-3FL (spc) ΔricF<sup>c</sup>* | This work |
| BD9210 | *rny-3FL (spc) ricT C161S* | This work |
| BD9211 | *rny-3FL (spc) ricT C198S* | This work |
| BD9212 | *rny-3FL (spc) ricT C167S* | This work |
| BD9217 | *P<sub>spac</sub>-ricT-3FL::thr (erm) rny::kan<sup>b</sup>* | This work |
| B9218 | *P<sub>spac</sub>-ricT-3FL::thr (erm) rny::kan<sup>b</sup> ricA::spc<sup>d</sup>* | This work |

<sup>a</sup>All strains were constructed in the IS75 background.
<sup>b</sup>These knockouts were obtained from the Bacillus Genetic Stock Center and were constructed by Koo et al (36).
<sup>c</sup>These deletions are markerless. Cassettes were removed using pDR244 (36).
<sup>d</sup>A gift from the late A. A. Neyfakh.

**Table S3.**
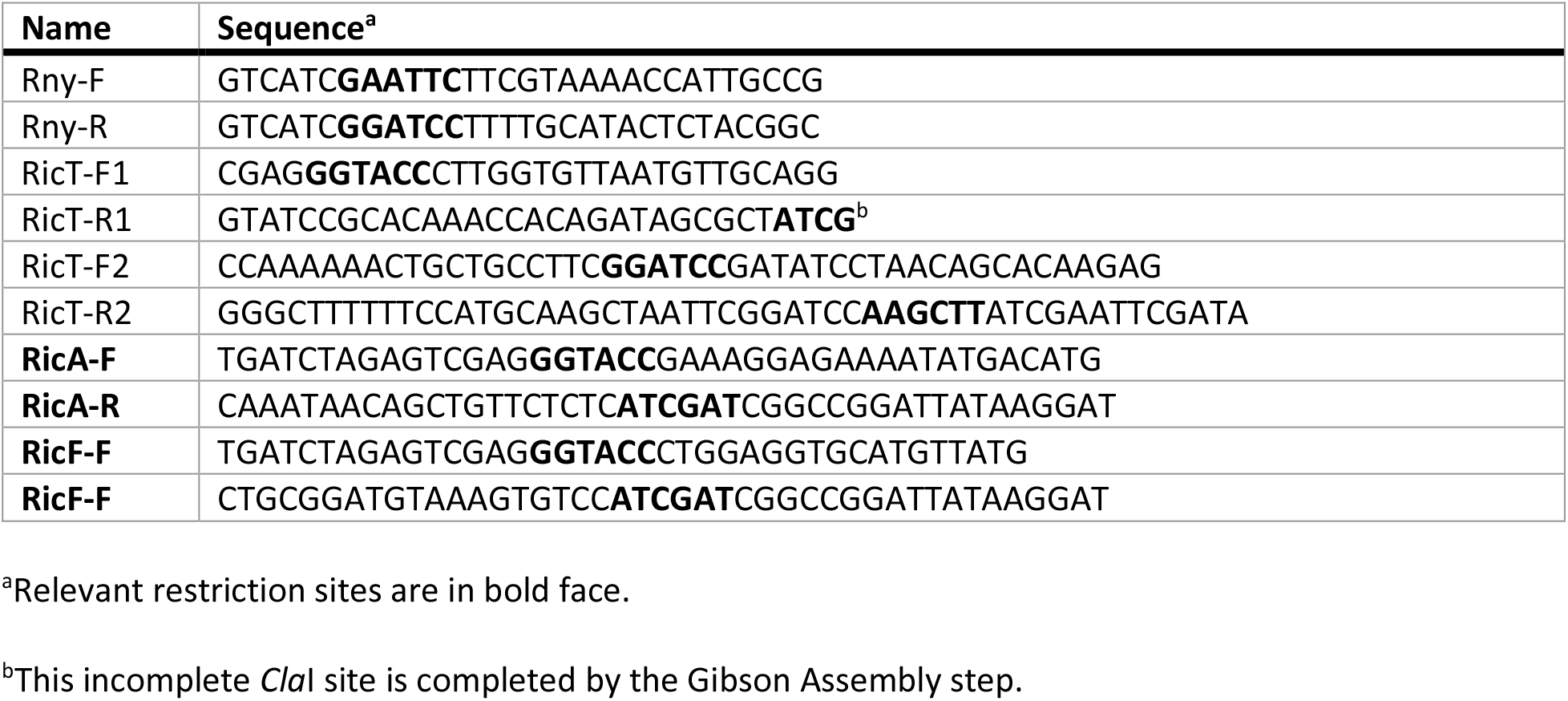
Oligonucleotides

